# LRScaf: Improving Draft Genomes Using Long Noisy Reads

**DOI:** 10.1101/374868

**Authors:** Mao Qin, Shigang Wu, Alun Li, Fengli Zhao, Hu Feng, Lulu Ding, Yuxiao Chang, Jue Ruan

**Affiliations:** Genome Analysis Laboratory of the Ministry of Agriculture, Agricultural Genomics Institute at Shenzhen, Chinese Academy of Agricultural Sciences, Shenzhen, Guangdong 518124, China.

**Keywords:** Assembly, Scaffolding, SMRT, ONT, Long Reads

## Abstract

**Background:** The advent of Third Generation Sequencing (TGS) technologies opens the door to improve genome assembly. Long reads are promised to enhance the quality of fragmental draft assemblies constructed from Next Generation Sequencing (NGS) technologies. To date, a few of algorithms, *i.e*., SSPACE-LongRead, OPERA-LG, SMIS, npScarf, DBG2OLC, Unicycler, and LINKS, have been released that are capable of improving draft assemblies. However, hybrid assembly on large genomes is still challenging.

**Results:** We develop a scalable and computationally efficient scaffolder, Long Reads Scaffolder (LRScaf), that is capable of boosting assembly contiguity to a large extent using long reads. In our experiment, our method significantly improves the contiguity of human draft assemblies, increasing the NG50 value of CHM1 from 127.5 Kb to 10.4 Mb using 20-fold coverage PacBio dataset and the NG50 value of NA12878 from 115.7 Kb to 17.4 Mb using 35-fold coverage Nanopore dataset. The run time for the scaffolding procedure using LRScaf is the shortest in all cases of our experiment. Compared with the run time of SSPACE-LongRead, LRScaf is faster 300 times for *S. cerevisiae* and 2,300 times for *D. melanogaster*. The peak RAM of LRScaf, by contrast, is more efficient than LINKS in our test. For the rice case, the peak RAM of LINKS (877.72 Gb) is about 196 times higher than LRScaf. For the experiment of human assembly, the peak RAM of LINKS is beyond the capacity of system memory (1 Tb) whereas LRScaf takes 20.28 and 41.20 Gb on CHM1 and NA12878 datasets.

**Conclusions:** The new method, LRScaf, yields the best or at least moderate contiguity and accuracy of scaffolds in the shortest run time compared with the state-of-the-art methods. Furthermore, it offers a new opportunity for the hybrid assembly of large genomes.

## Background

With the advent of Next Generation Sequencing (NGS) technologies, the genomics community has made significant contributions to *de novo* assembling genomes. Despite that many studies and tools are aimed at reconstructing NGS data into complete *de novo* assemblies of genomes, this goal is difficult to achieve because of intrinsic limitation of NGS data, *i.e*., read lengths are shorter than most of the repetitive sequences [1]. The existence of repeats makes it difficult to reconstruct complete genomes instead of generating a large set of contiguous sequences (contigs) even when the sequencing coverage is high [2]. Thus, attention is focused on the so-called genomic scaffolding procedure, which aims at reducing the number of contigs by using fragments of moderate lengths whose ends are sequenced (double-barreled data) [3,4]. Nevertheless, major genomic regions still hinder genomic assemblies because of, primarily, large-size repeat and low coverage. In response, Third Generation Sequencing (TGS) technologies have been developed. TGS sheds light on different alternatives to solve genome assembly problems by offering very long reads, *e.g*., the Single Molecule Real Time (SMRT) sequencing technology of Pacific Biosciences^®^ (**PacBio**)delivers read lengths of up to 50 Kb [5] and the nanopore sequencing technology of Oxford Nanopore Technologies^®^ (ONT) delivers read lengths which are greater than 800 Kb [6]. These long reads suffer from high sequencing error rates, however, which necessitates high coverage during the genome assembly [7]. In addition, TGS technologies have a higher cost per base than NGS methods. Consequently, long reads are more commonly used for scaffolding draft assemblies generated from NGS data than for *de novo* assembly [8].

The process of genome assembly is typically divided into two major steps. The first step is to piece overlapping reads together into contigs which is commonly done using the *de Bruijn* or overlap graph [1]. The second step is to assemble scaffolds, consisting of ordered sequences of oriented contigs with estimated distances between them. Scaffolding, which was first introduced by Huson [3], is a critical part of the genome assembly process, especially for NGS data. Yet, scaffolding is a research area that remains largely open because of the NP-hard complexity [9]. By using paired-end and/or mate-pair reads linking information, a number of standalone scaffolders, *e.g*. Bambus [4], MIP [10], Opera [11], SCARPA [12], SOPRA [13], SSPACE [14], BESST [15], and BOSS [16], have been developed. Nevertheless, a recent comprehensive evaluation showed that scaffolding was still computationally intractable and required better quality large insert-size pair read libraries than presently available [17]. As TGS technologies are likely to offer longer reads than the lengths of the most common repeats, these technologies are capable of drastically reducing and solving the complexity caused by repeats. Considered the pros and cons of NGS and TGS data, a hybrid assembly approach that assembled draft genomes using TGS data was proposed [18]. The core strategy of this approach is: 1) long reads are mapped onto the contigs using a long-read mapper (e.g. BLASR [19] or minimap [20,21]); 2) examining alignment information, long reads that span more than one contig are identified and their linking relationship is stored in a data structure; 3) the last step is to clean up the structure by removing redundant and error-prone links, calculate distances between contigs, and build scaffolds using links information.

Based on the hybrid assembly strategy, AHA [18] was the first standalone hybrid scaffolder and was part of the SMRT analysis software suite. As AHA was designed for small genomes and had limitations on the input data, it was not suitable for large genomes. To ensure that scaffolds were as contiguity as possible, AHA performed 6 iterations by default, thus increasing the run time. SSPACE-LongRead [22] produced the final scaffolds in a single iteration and, therefore, had a significantly shorter run time than AHA. Nevertheless, SSPACE-LongRead had somewhat lower assembly accuracy than AHA. Despite being designed for large eukaryotic genomes, SSPACE-LongRead was unpractical because of its intensive run time. LINKS [23] opened a new door to build linking information between contigs. The algorithm used the long interval nucleotide K-mer without computational alignment and reads correction step, but its memory usage was a concern. OPERA-LG [24] provided an exact algorithm for large and repeat-rich genomes. Its main limitation was that it required significant mate-pair information to constrain the scaffold graph and report an optimized result. OPERA-LG was not directly designed for TGS data, and to construct scaffold edges and link contigs together into scaffolds, OPERA-LG needed to be modified by simulated and grouped mate-pair relationship information from long reads. Recent studies, such as SMIS (Available from http://www.sanger.ac.uk/science/tools/smis), npScarf [25], DBG2OLC [26] and Unicycler [27], have been reported based on the hybrid assembly strategy. However, these tools have not been thoroughly assessed for different genome sizes, especially large genomes.

Here we present a Long Reads Scaffolder (LRScaf) to improve draft genomes using TGS data. The input to LRScaf is given by a set of contigs and their alignments over SMRT or ONT long reads. We compare our method with the state-of-the-art tools on real and synthetic datasets. All the methods tested improve the contiguity of pre-assembled genomes. Our method yields the best assembly metrics and contiguity for pre-assembled genomes of *E. coli, S. cerevisiae, D. melanogaster*, and *H. sapiens*. More importantly, however, our method consistently returns the most accurate scaffolds and has the shortest run time. Especially, LRScaf significantly improves the contiguity of human draft assemblies, increasing the NG50 value of CHM1 from 127.5 Kb to 10.4 Mb using 20-fold coverage PacBio dataset and the NG50 value of NA12878 from 115.7 Kb to 17.4 Mb using 35-fold coverage Nanopore dataset. We thus show that LRScaf is a valuable tool for improving draft assemblies in a cost-effective way.

## Results and discussion

We performed in-depth analysis on five species, *i.e., E. coli, S. cerevisiae, D. melanogaster, O. sativa*, and *H. sapiens*, to test and compare the performance of LRScaf with that of SMIS, npScarf, DBG2OLC, Unicycler, SSPACE-LongRead, LINKS, and OPERA-LG. The details of datasets are provided in Table 1 and in the Methods section. The NGS datasets for *E. coli* and *S. cerevisiae* are real with 600 and 105 -fold coverages respectively, where the NGS datasets for *D. melanogaster* and *O. sativa* were synthesized using pIRS [28] with 50-fold coverage. The real reads for the two small genomes (*E. coli* and *S. cerevisiae*) were first cleaned and then used to construct draft assemblies using SOAPdenovo2 [29] and SPAdes [30]. The synthetic reads for the two large genomes (*D. melanogaster* and *O. sativa*) were directly used to build draft assemblies using SOAPdenovo2. The draft assemblies of two human lines CHM1 [31] and NA12878 [32] were used to test the performances of all scaffolders for large genomes using the PacBio and Nanopore datasets. The statistics of draft assemblies are shown in Table 2. Results and assembly metrics obtained after the scaffolding procedure are displayed in Tables 3 and 4.

**Table 1.**
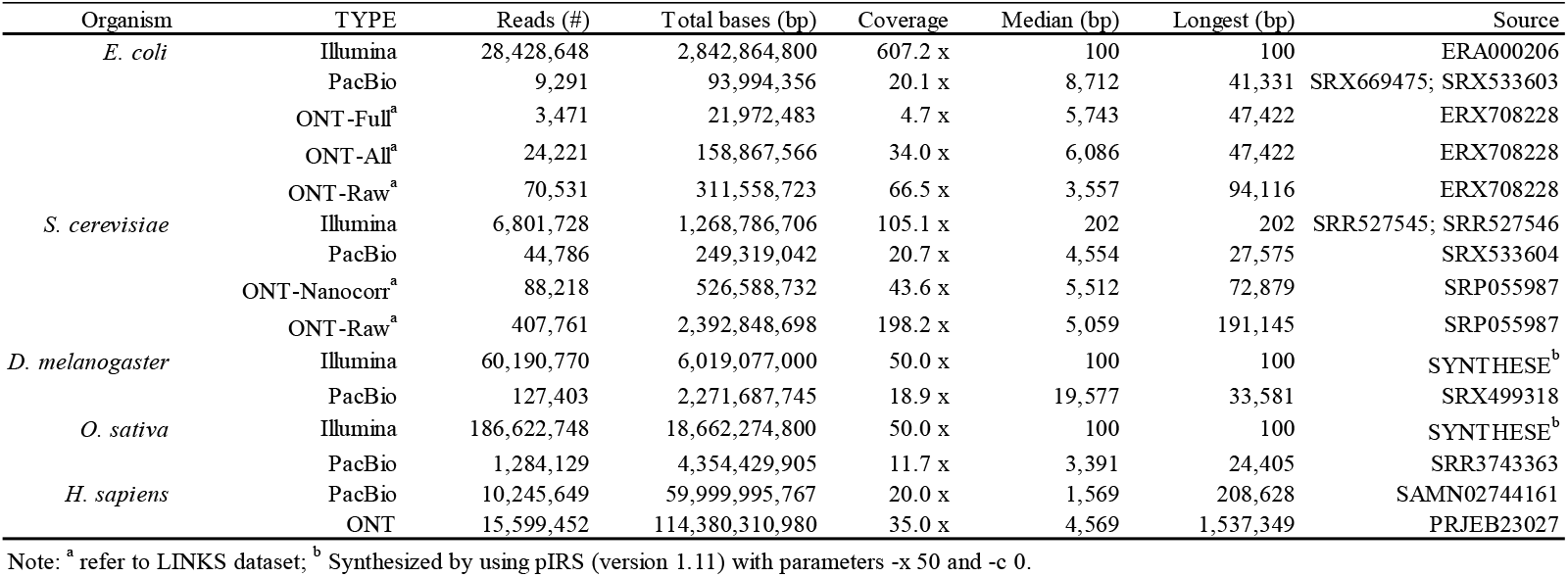
Descriptive statistics of datasets used for the comparative study.

**Table 2.** Draft assembly statistics for *E. coli*., *S. cerevisiae*, *D. melanogaster*, *O. sativa*, and *H. sapiens*.

| Organism | Source | Reference Length (bp) | Chr. | Assembled Length (bp) | Fraction | Contigs (#) | N00 (bp) | N50 (bp) | N100 (bp) |
| --- | --- | --- | --- | --- | --- | --- | --- | --- | --- |
| *E. coli* | SOAPdenovo2 | 4,681,865 | 1 | 4,598,322 | 0.98 | 728 | 164,235 | 40,009 | 52 |
|  | SPAdes |  |  | 4,579,398 | 0.98 | 242 | 264,985 | 133,189 | 56 |
|  | ABYSS<sup>a</sup> |  |  | 5,160,631 | 1.10 | 69 | 358,719 | 177,636 | 493 |
| *S. cerevisiae* | SOAPdenovo2 | 12,071,326 | 16 | 12,063,232 | 0.99 | 6,961 | 146,672 | 19,567 | 60 |
|  | SPAdes |  |  | 11,754,316 | 0.97 | 2,254 | 451,383 | 107,906 | 56 |
|  | Celera Assembly<sup>a</sup> |  |  | 14,910,895 | 1.24 | 6,953 | 257,346 | 49,258 | 64 |
| *D. melanogaster* | SOAPdenovo2 | 120,381,546 | 6 | 118,065,428 | 0.98 | 45,480 | 902,599 | 111,033 | 62 |
| *O. sativa* | SOAPdenovo2 | 373,245,519 | 12 | 346,168,844 | 0.93 | 257,801 | 147,060 | 18,977 | 3 |
| *H. sapiens* | SRPRISM+ARGO<sup>b</sup> | 2,996,426,293 | 23 | 2,781,084,252 | 0.93 | 40,906 | 1,009,096 | 140,502 | 199 |
|  | DISCOVAR<sup>c</sup> |  |  | 3,068,057,564 | 1.02 | 858,918 | 1,380,479 | 179,783 | 201 |

Note: <sup>a</sup> refers to LINKS; <sup>b</sup> refers to [31]; <sup>c</sup> refers to [32].

**Table 3.**
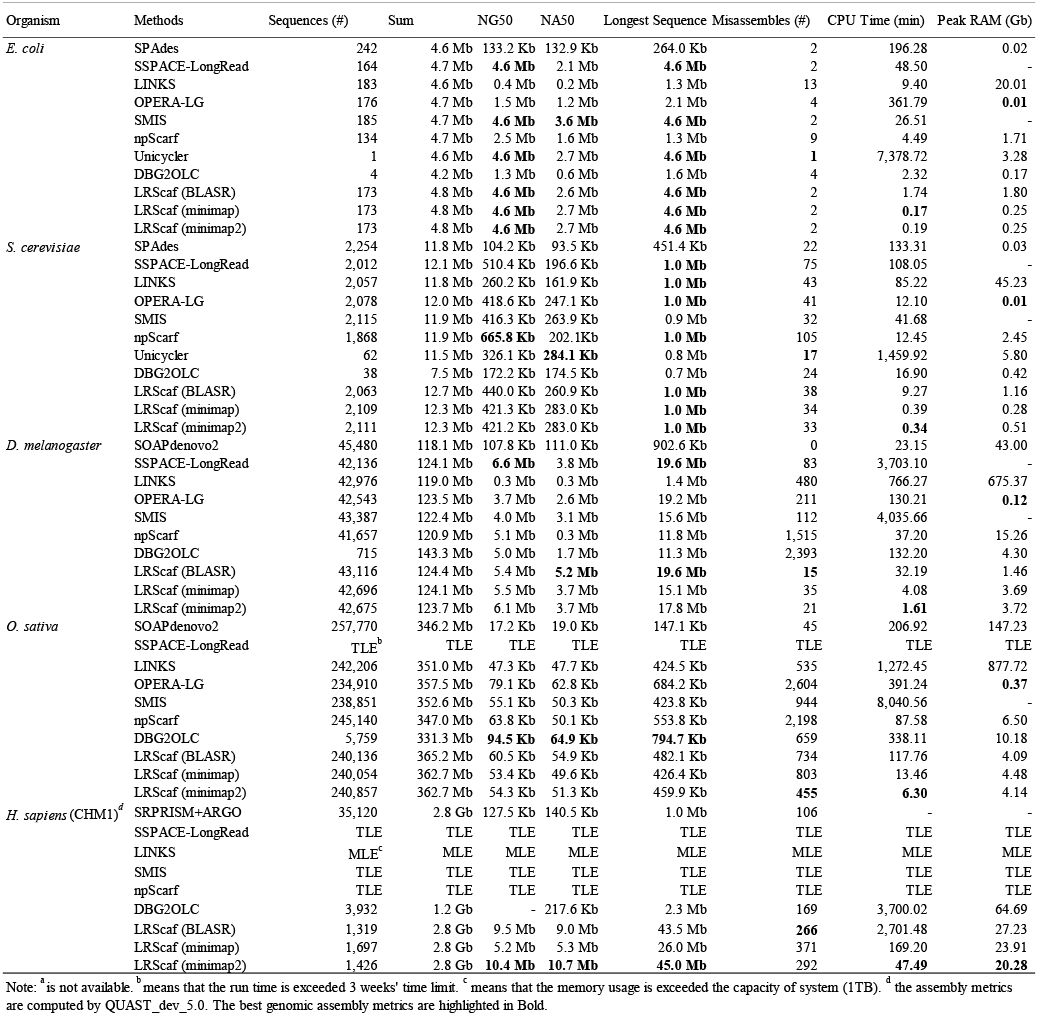
The performances of scaffolders tested for *E. coli*, S. *cerevisiae*, *D. melanoçaster*, *O. sativa*, and *H. sapiens* using PacBio long reads.

**Table 4.**
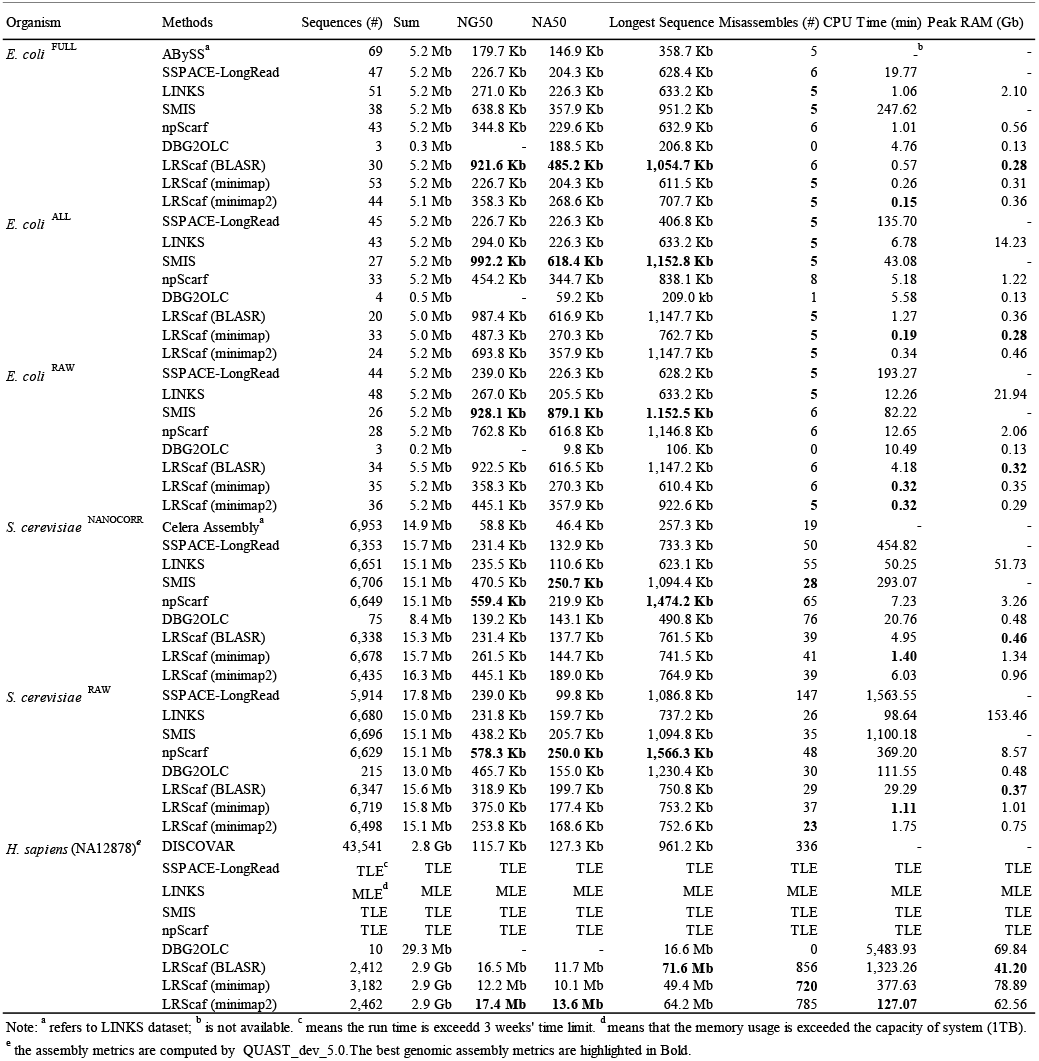
The performances of scaffolders tested for *E. coli, S. cerevisiae*, and *H. sapiens* using ONT long reads.

### Draft genome assemblies

We used SPAdes to construct draft assemblies for two small genomes *(E. coli* and *S. cerevisiae)*with the “careful” parameter option. Draft assemblies for these two small genomes were also constructed using SOAPdenovo2. In addition, SOAPdenovo2 was used to construct draft assemblies for *D. melanogaster* and *O. sativa*, whose synthetic reads were available. We used the optimal k-mer values for the draft assemblies constructed by SOAPdenovo2 with 51 (*E. coli*), 59 (*S. cerevisiae*), 61 (*D. melanogaster*), and 73 (*O. sativa*). These values were selected based on assembled genome size, number of contigs, and genome contiguity.

The statistics of draft assemblies for *E. coli*, *S. cerevisiae*, *D. melanogaster*, *O. sativa*, and *H. sapiens* are shown in Table 2. For *E. coli*, the draft-genome size obtained using SOAPdenovo2 is 4.6 Mb distributed over 728 contigs, yielding an assembled genome fraction of 98 % and an N50 value of 40.0 Kb. SPAdes yields a draft-assemblies size of 4.6 Mb with 242 contigs, a genome fraction of 98 %, and an N50 value of 133.2 Kb. The draft-assemblies size generated by ABySS is 5.2 Mb with 69 contigs, a genome fraction of 110 %, and an N50 value of 177.6 Kb. For *S. cerevisiae*, the draft-genome size obtained using SOAPdenovo2 is 12.1 Mb with 6,961 contigs, providing a genome fraction of 99 % and an N50 value of 20.0 Kb. SPAdes generates a draft-assemblies size of 11.8 Mb with 2,254 contigs, yielding a genome fraction of 97 % and an N50 value of 107.9 Kb. The draft-assemblies size constructed by Celera Assembly is 15.0 Mb with 6,953 contigs, a genome fraction of 124 %, and an N50 value of 49.2 Kb. For *D. melanogaster*, the draft-assemblies size constructed by SOAPdenovo2 is 118.1 Mb with 45,480 contigs, a genomic fraction of 98 %, and an N50 value of 111.0 Kb. The draft-genome size for *O. sativa* is 346.2 Mb and it contains 257,801 contigs, yielding a genomic fraction of 92 % and an N50 value of 19.0 Kb. The size of the draft genome of CHM1 is 2.8 Gb distributed over 40,906 contigs, and it has a genomic fraction of 93 % and an N50 value of 140.0 Kb where the draft-assemblies size of NA12878 is 3.1 Gb with 858,918 contigs, yielding a genome fraction of 102 % and an N50 value of 179.8 Kb.

The depth of coverage is an important factor in *de novo* genome assembly. The genome contiguity and completeness obtained are not only determined by the depth of coverage, however, but also by the method’s ability to overcome complex genome structures, *e.g*. repetitive regions. *E. coli* is the smallest genome and has the highest coverage (more than 600-fold of NGS reads) among the genomes included in this study. However, the assembly contiguity is still fragmental. As the genome gets larger and more complex, draft assemblies become increasingly fragmental unless auxiliary technologies are included in the assembly process. Consequently, the inclusion of large insert-size mate-pair libraries, Hi-C [33], optical-mapping data [34] and long reads is important to overcome large repeats and to assist the scaffolding procedure.

### Scaffolding on SMRT long reads

In this study, we used long reads of SMRT datasets for *E. coli, S. cerevisiae, D. melanogaster, O. sativa*, and *H. sapiens* to assess the performances of seven state-of-the-art scaffolders (*i.e*., SSPACE-LongRead, LINKS, OPERA-LG, SMIS, npScarf, Unicycler, and DBG2OLC) and our LRScaf (See Table 1). The median lengths of SMRT long reads for 5 organisms are 8.7 Kb, 4.6 Kb, 19.6 Kb, 3.4 Kb, and 1.6 Kb, respectively. And the longest reads are 41.3 Kb, 27.6 Kb, 33.6 Kb, 24.4 Kb, and 208.6 Kb, respectively. The coverages of SMRT long reads are 20.1-fold (*E. coli*), 20.7-fold (S. *cerevisiae*), 18.9-fold (*D. melanogaster*), 11.7-fold (O. *sativa*), and 20.0-fold (*H. sapiens*). The distributions of read length show that the SMRT long reads approximate normal distributions (See Suppl. Fig. 1). The SMRT long reads of *D. melanogaster* were filtered for the FALCON assembler [35], which resulted in an increased average read length. QUAST [36] was used to assess draft assemblies after the scaffolding procedure. The released version 4.5 of QUAST was failed to assess human assemblies, and, therefore, we used the dev-5.0 version to evaluate the corresponding assembly metrics.

All scaffolders reduce the number of contigs and improve assemblies contiguity (See Table 3 and Suppl. Tables 1 and 2). Whereas SSPACE-LongRead, SMIS, Unicycler, and LRScaf reconstruct the genome for *E. coli* into a complete single chromosome, LINKS, OPERA-LG, npScarf, and DBG2OLC fail to do that. In addition, Unicycler significantly reduces the numbers of contigs. For the 1, 5, and 10 -fold coverages, the performances of scaffolders tested show similar results on the 20-fold coverage where the assemblies contiguity of SSPACE-LongRead, SMIS, Unicycler, and LRScaf are better than that of LINKS, OPERA-LG, npScarf, and DBG2OLC (See Suppl. Tables 1 and 2). For *S. cerevisiae*, the npScarf method yields the best NG50 value (665.8 Kb) and Unicycler generates the best NA50 value (284.1 Kb). SSPACE-LongRead, LINKS, OPERA-LG, npScarf, and LRScaf yield the longest sequence (1.0 Mb). For the 1, 5, and 10 -fold coverages, SSPACE-LongRead yields the best assemblies contiguity (NG50) in 5 out of 6 cases and OPERA-LG, npScarf, and LRScaf yield the best NG50 in 1 out of 6 cases (See Suppl. Table 1). Based on draft assemblies generated by SOAPdenovo2 using 20-fold coverage, SSPACE-LongRead and LRScaf yield the best NG50 and NA50 value respectively and generate the longest sequence (See Suppl. Table 2). For *D. melanogaster*, SSPACE-LongRead yields the best NG50 value (6.6 Mb) and LRScaf with BLASR produces the best NA50 value (5.2 Mb). SSPACE-LongRead and LRScaf construct the longest sequence of 19.6 Mb. For *O. sativa*, DBG2OLC significantly reduces the number of sequences and produces the best NG50 value (94.5 Kb) and NA50 value (64.9 Kb), and the longest sequence (794.7 Kb). SSPACE-LongRead is excluded from this assessment because it exceeds the 3 weeks’ run time limit. For *H. sapiens* CHM1, LRScaf with minimap2 yields the best NG50 value (10.4 Mb) and NA50 value (10.7 Mb,), and the longest sequence (45.0 Mb). The run time of SSPACE-LongRead, SMIS, and npScarf exceeds the time limit, and LINKS exceeds our system’s memory capacity of 1 Tb. Thus, these scaffolders are excluded from the test on the *H. sapiens* CHM1 genome. As evident from our experiments, the run time and the memory usage for these scaffolders become significant concerns for the large and complex genomes. DBG2OLC is recommended to use SparseAssembler (Available from: https://github.com/yechengxi/SparseAssembler) to construct draft assemblies for hybrid assembly. This might be the reason for the assembly genome size generated by DBG2OLC is smaller than what the other scaffolders yield, especially for the *H. sapiens*. To summarize, LRScaf yields the best or, at least, moderate assembly metrics when compared with other scaffolders on SMRT long reads.

### Scaffolding on ONT long reads

We used the ONT long reads datasets for *E. coli*, *S. cerevisiae*, and *H. sapiens* to assess the performances of scaffolders tested (See Table 4). Because of lack of NGS data, OPERA-LG and Unicycler were excluded from this assessment. For the two small genomes, the ONT long-reads datasets were published in LINKS, including 3 of *E. coli* (FULL, ALL and RAW datasets with 4.7, 34.0, and 66.5 -fold coverages, respectively) and 2 of *S. cerevisiae* (NANOCORR and RAW datasets with 43.6 and 198.2 -fold coverages). We used the *H. sapiens* NA12878 dataset with 35.0-fold coverage as the large genome for this test. The best median and longest length of reads are 6.1 Kb and 1.5 Mb respectively (See Table 1). The distributions of read length show that ONT long reads approximate bimodal distributions with a long tail (See Suppl. Fig. 2). The median length of ONT reads is approximately equal to that of SMRT, but the longest length of ONT reads is significantly longer than that of SMRT datasets. QUAST (Version 4.5) was used to assess draft assemblies and scaffolded assemblies for *E. coli* and *S. cerevisiae*. And QUAST (Dev-5.0 version) was used to evaluate the corresponding assembly metrics for *H. sapiens*.

All scaffolders decrease the number of contigs and improve genome contiguity (See Table 4). The number of contigs for the *E. coli* draft assemblies is 69 with an NG50 value of 179.7 Kb. For the FULL dataset, LRScaf with BLASR yields the best NG50 value (921.6 Kb) and NA50 value (485.2 Kb), and the longest sequence (1.1 Mb). SMIS generates the best NG50 value (992.2 Kb) and NA50 value (618.4 Kb), and the longest sequence (1.2 Mb) for the ALL dataset where LRScaf with BLASR yields similar performance (NG50: 922.5 Kb, NA50: 616.5 Kb, and the longest sequence 1.1 Mb). Whereas SMIS produces the best numbers for the RAW dataset (NG50: 928.1 Kb, NA50: 879.1 Kb, and the longest sequence 1.2 Mb), LRScaf with BLASR yields very similar metrics. For *S. cerevisiae*, the number of contigs for the draft assemblies is 6,953 with an NG50 value of 58.8 Kb. For the NANOCORR dataset, the npScarf method yields the best NG50 value and the longest sequence (559.4 Kb and 1.5 Mb, respectively), and SMIS produces the best NA50 value (250.7 Kb). The npScarf scaffolder also produces the best metrics (NG50: 578.3 Kb, NA50: 250.0 Kb, and the longest sequencing 1.6 Mb) for the RAW dataset. For the NA12878 dataset, LRScaf with minimap2 significantly improves the contiguity of the draft assemblies and yields the best NG50 and NA50 values (17.4 Mb and 13.6 Mb, respectively). LRScaf with BLASR produces the longest sequence (71.6 Mb). All the other scaffolders are similar to the assessment using the PacBio dataset and exceed either the time limit (3 weeks) or the memory capacity of system (1 Tb). In addition, DBG2OLC is not successful to scaffold draft assemblies generated by DISCOVAR. This is as expected where DBG2OLC is recommended to use SparseAssembler as its NGS Assembler for hybrid assembly. Compared with the results obtained using the SMRT datasets, none of the scaffolders could assemble *E. coli* into a single chromosome and the contiguity of *S. cerevisiae* is more fragmented. Although all scaffolders show certain improvement in our experiment, the application of the ONT data is still challenging. A recent study showed that the NA12878 genome was assembled with an NG50 value of about 6.5 Mb using pure 35-fold ONT data [6]. Our experiments, however, show that it is possible to significantly improve assembly contiguity to 17.4 Mb where it is similar to the PacBio human case. To summarize, LRScaf yields either the best or similar assembly metrics using long reads of ONT compared with the other scaffolders.

### Computational performance and accuracy analysis

The assembly metrics are undoubtedly the most concerning matters to biologists and bioinformaticians. Nevertheless, from a practical point of view, the run time limits software applications. SSPACE-LongRead and OPERA-LG use BLASR as their default TGS mapper for construction of joints between contigs. The npScarf software uses BWA [37] as its default mapper. LINKS, SMIS, Unicycler, and DBG2OLC use its built-in algorithms to build joints between contigs. To enable a direct comparison with SSPACE-LongRead and OPERA-LG, our LRScaf supports BLASR. Nevertheless, it also supports a faster TGS mapper minimap (Versions 1 and 2), which enables a significant reduction for the total run time of the scaffolding procedure. LRScaf is the fastest scaffolder for all the cases using SMRT long reads. LRScaf reduces the run time more than 300 times compared with SSPACE-LongRead and more than 3,900 times compared with Unicycler for *S. cerevisiae* (See Table 3). As the genome gets larger, the advantage of shorter run time becomes more important. In *D. melanogaster*, LRScaf is 2,300 times faster than SSPACE-LongRead and 2,550 times faster than SMIS. In *O. sativa*, LRScaf is 1,276 times faster than SIMS. We have no number on how much LRScaf is faster than SSPACE-LongRead because the latter exceeds the time limit (3 weeks). For *H. sapiens*, SSPACE-LongRead, SMIS, and npScarf exceed the time limit (3 weeks). For the ONT datasets, LRScaf is also the fastest scaffolder. LRScaf is more than 131 times faster than SSPACE-LongRead on the FULL dataset for *E. coli*. As the dataset grows larger, the advantage becomes more significant. LRScaf is 714 times faster on the ALL dataset for *E. coli*, 603 times faster on the RAW dataset for *E. coli*, and 1,408 times on the RAW dataset for *S. cerevisiae* than

SSPACE-LongRead. LINKS skips the all-to-all alignment step and is faster than SSPACE-LongRead in all cases. Nevertheless, the memory usage of LINKS is of concern and it might be alleviated by further improvement of the data-structure. Although the peak RAM usage for LRScaf is higher than that of OPERA-LG on small genomes, our experiments show that the memory usage of LRScaf is practical even for large and complex genomes where the peak RAM for LRScaf is not over 30 Gb on CHM1 PacBio dataset and 80 Gb on NA12878 ONT dataset.

Reducing the number of misassemblies is important because misassemblies are likely misinterpreted as true genetic variations [38,39]. For the SMRT datasets, SSPACE-LongRead and LRScaf yield the fewest number of misassemblies (1) among the scaffolders based on draft assemblies for *E. coli* generated by SOAPdenovo2 (See Suppl. Table 2). Unicycler produces the fewest number of misassemblies for *E. coli* (1) and *S. cerevisiae* (17) based on draft assemblies constructed by SPAdes where LINKS and npScarf yields the maximum number of misassemblies for *E. coli* (13) and *S. cerevisiae* (105) respectively (See Table 3). LRScaf yields the fewest number of misassemblies for *D. melanogaster* (15) and *O. sativa* (455) where DBG2OLC and OPERA-LG produce the maximum number of misassemblies for *D. melanogaster* (2,393) and *O. sativa* (2,604) respectively (See Table 3). For *H. sapiens*, we have no number on how many the number of misassemblies for the other scaffolders because all of them are failed to scaffold the draft assemblies. For the ONT datasets, the draft assemblies for *E. coli, S. cerevisiae*, and *H. sapiens* contain 5, 19, and 336 misassembled contigs, respectively, and none of the scaffolders significantly increases the number of misassemblies (See Table 4). LRScaf with minimap2 outputs the fewest number of misassemblies on the *E. coli, S. cerevisiae* (RAW data). LRScaf with minimap outputs the fewest number of misassemblies on *H. sapiens*. SMIS yields the fewest number of misassemblies for the *S. cerevisiae* NANOCORR dataset. SSPACE-LongRead yields the maximum number of misassemblies (147) on the RAW dataset for *S. cerevisiae*. In summary, LRScaf introduces a new strategy for keeping valid alignments (See Methods section) and produces fewer misassemblies than most of the other scaffolders. Moreover, LRScaf with minimap2 significantly reduces the run time of scaffolding procedure without increasing the number of misassemblies. Based on the SMRT and ONT performances, we recommend that LRScaf is used with BLASR on small genomes and with minimap on large genomes.

## Conclusion

In this work, we present a novel program for scaffolding draft assemblies using noisy TGS long reads information and compare our algorithm with the previous methods. The majority of the draft assemblies constructed using NGS data is fragmented and influenced by repeats. The disadvantage of long reads is that they contain significantly more errors than first- and second-generation sequencing technologies. Nevertheless, we successfully use long reads to build links between contigs, overcome repetitive regions, and improve genome contiguity. We propose a new strategy to filter inaccurate alignments so that these false alignments do not propagate through the scaffolding process. For the assessments on SMRT long-read datasets covering 5 organisms, our method shows significant improvements over the state-of-the-art scaffolders. The primary benefits of LRScaf over these scaffolders are that it yields the fewer number of misassemblies and reduces the run time, yet it retains the best or, at least, average assembly metrics. These improvements are especially useful for large and complex genomes. For the assessments on ONT long-read datasets for 3 organisms, our method shows significant improvements over the previous algorithms. Our method keeps the best or, at least, average assembly metrics and the shortest run time. In addition, our method has the fewest number of misassemblies in most of the cases. As studied genomes keep getting larger and more complex, the run time and the memory usage for the analysis software are becoming increasingly important to biologists and bioinformaticians. Our method is designed with reduction of the run time and the memory usage in mind and is, thus, much faster than other scaffolders and requires only moderate memory usage. Identification of misassembled contigs is also important, however, because any misassembled sequences are propagated into the next step during biological analysis. Most state-of-the-art scaffolders lack functions for identification of misassembled contigs. In addition, misassemblies might be introduced during the scaffolding procedure. Consequently, to limit the number of misassembled scaffolds, our method incorporates a validation algorithm that checks the links information between contigs. As checking and correcting misassemblies from draft assemblies is important, we are planning to use long read information to achieve and integrate these functions in a future version of LRScaf.

In the past decade, worldwide collaboration has led to several projects, aiming at improving the understanding of species biology and evolution. Examples of such projects are the i5k [40], which provides the genomes of 5,000 species of insects, and the Bird 10,000 Genomes (B10K) [41]. However, a substantial fraction of genomes with short contiguity hinder downstream analysis. Our result shows that TGS data is capable of effectively improving draft assemblies and LRScaf is a valuable tool for improving draft assemblies in a cost-effective way.

## Methods

### Alignment of TGS long reads

LRScaf was designed to separate the mapping and scaffolding procedures. Hence, during the mapping procedure, we set the number of processes to 48 and kept the default values for all other parameters using BLASR and minimap (Version 1 and 2). LRScaf supports the default alignment format of these mappers.

### Validating alignment

The high error rate is a serious disadvantage of TGS long reads. Thus, a large fraction of the alignments is incorrect and needs to be filtered out. We developed a validation model to validate each alignment (See Figure 1). The model partitioned each long read into three regions (R1, R2, and R3) separated by two points (P1 and P2). Considered the alignment start (S) and end (E) loci in the contig, there were six different combination sets in *R*, *i.e*., ***R* ∈ {(*S in R*1, *E in R*1), (*S in R*1, *E in R*2), (*S in R*1, *E in R*3), (*S in R*2, *E in R*2), (*S in R*2, *E in R*3), (*S in R*3, *E in R*3)}**. We also defined the distal length of a contig to the start or end alignment loci as the over-hang length of the contig. Taken both the alignment region and the over-hang length into account, the valid alignment satisfied: 1) (S in R1, E in R1) with the right over-hang length not exceeding the constraints; 2) (S in R1, E in R2) with the right over-hang length not exceeding the constraints; 3) (S in R2, E in R2) with the two end over-hang length not exceeding the constraints; 4) (S in R2, E in R3) with the left over-hang length not exceeding the constraints; 5) (S in R3, E in R3) with the left over-hang length not exceeding the constraints. An alignment was filtered out if a long read was entirely covered by a contig (S in R1, E in R3), *i.e*., the contig contained the long read. After this procedure, the remaining alignments were considered to be valid for the scaffolding procedure.

**Figure 1.**
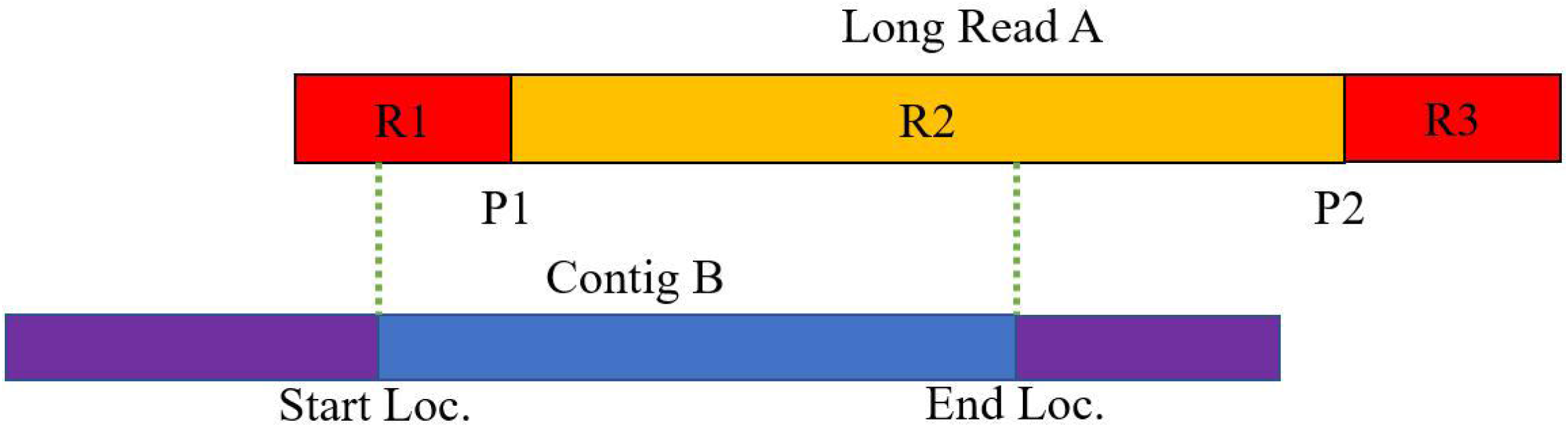
A validating model of alignment. The P1 and P2 are the two points for breaking a long read into 3 regions (R1, R2, and R3).

### Repeat identification

Repetitive sequences complicate the genome assembly. Thus, such sequences were masked in our approach. First, based on the uniform coverage of TGS data, we identified and removed repeats by the coverage of reads. In the calculation of reads coverage, long reads that covered the entire contig were counted. Then we computed the mean coverage and the standard deviation among the set of contigs. Any contig coverage that was larger than the threshold coverage, which was set to *μ_cov_* + 3 × *s.d._cov_*, was considered to be a repeat and the corresponding contig was removed from the next step of the analysis.

### Constructing links and edges

A long read may have multiple mappings because of repeats and high sequencing error rate. Figure 2 describes how links are built between contigs from the validated alignments. This process had two constraints on orientation and distance. Four strand combination sets *S* were used between contigs to constrain orientation, *i.e*., ***S* ∈ {*s*_1_: (+,+), *s*_2_: (+, −), *s*_3_: (−, +), *s*_4_: (−,−)}**. We defined the orientation between contigs as *O*(*c_i_*,*c_j_*) = *max* (*s*). The probability that the internal distance *e* between two contigs lies outside the range [*μ_is_* − 3 × *σ_is_*] was less than 5%, because *e* approximately follows a normal distribution *N*{*μ_is_,σ_is_*). If *e* lay outside the range [*μ_is_* − 3 × *σ_is_,μ_is_* + 3 × *σ_is_*], it was considered to be abnormal and the linking information was removed. Any long reads linking a contig to itself at different loci were also removed. After validating two constraints on links between contigs, we introduced an edge to represent a bundle of links that jointed two contigs using quadruple parameters 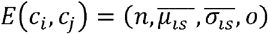. Here, *n* was the number of remaining links considered as the weight of the edge, 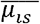 was the mean internal distance for the remaining links, 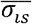 was the standard deviation of the internal distances for the remaining links, and *o* was the orientation strand between contigs.

**Figure 2.**
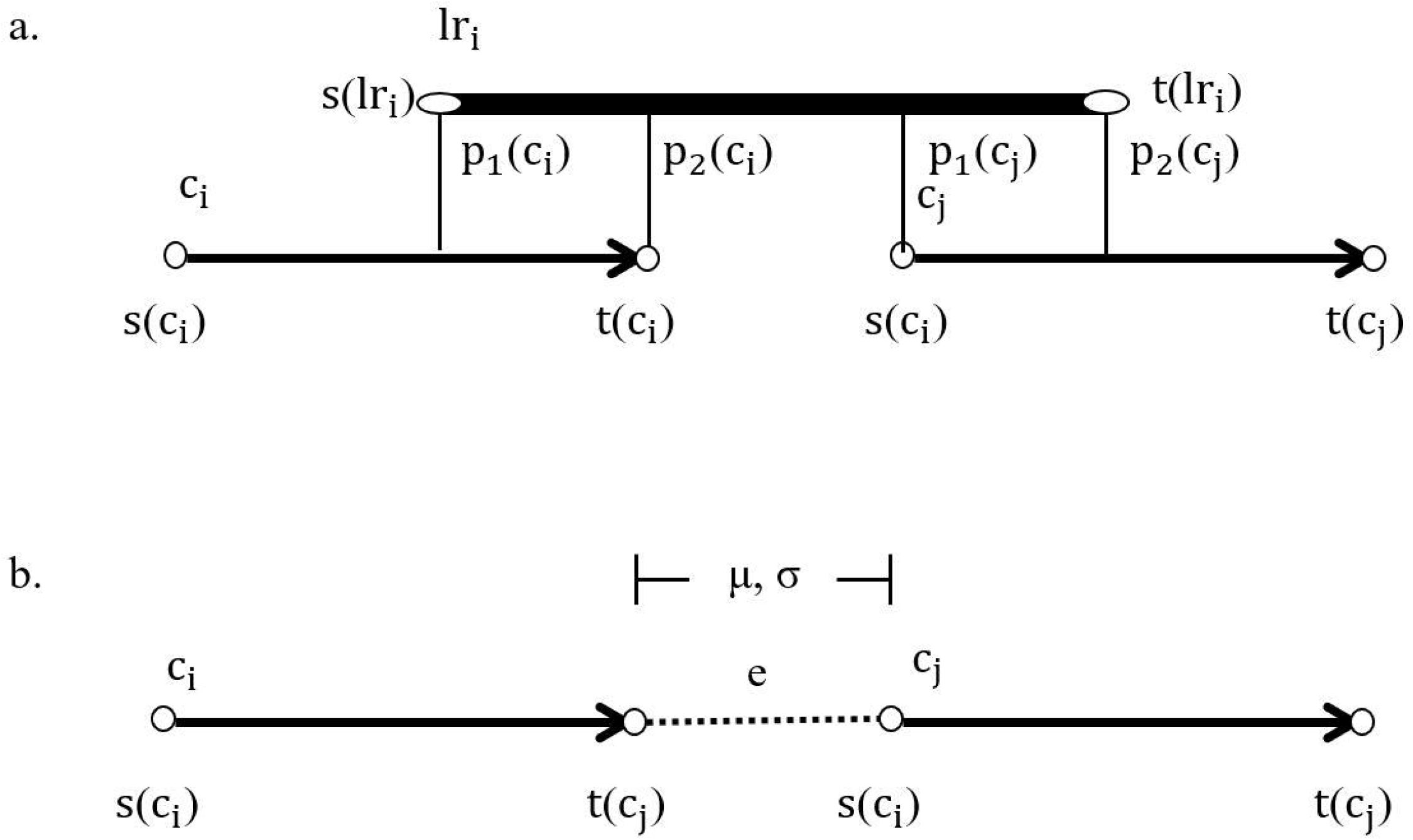
The construction of link using a long read *lri* and two contigs *c_i_* and *c_j_*. a) a basic schematic for a long read building link between contigs; b) the distance distribution of links.

### Graph construction and simplification

In this step, LRScaf constructed a scaffold graph *G*(*V, E*) similar to the string graph formulation. The vertex set *V* represented the end of the contigs and the edge set *E* represented the linkage implied by long reads between ends of two contigs with weight and orientation function assigned to each edge. The ends of each contig were annotated by their ID with a forward strand (+). Used this node concept, there were 4 types of edges in the graph, *i.e*., (+, +) joining the forward strands of both contigs, (+, −) joining the forward strand of the first contig with the reversed strand of the second contig, (−, +) joining the reversed strand of the first contig with the forward strand of the second contig, and (−, −) joining the reversed strands of both contigs. After the edges-construction step, we accounted for the majority of the sequencing errors by removing all the edges that had a lower number of long reads than the threshold value. Once the edges were cleaned and filtered, we constructed an assembly graph *G*. We only added an edge to *G* if neither of the two nodes comprising the edge was present in *G*. In some cases, *G* contained some edges of transitive reduction, error-prone and tips. Thus, such edges were deleted and we got the final scaffold graph which we used for further analysis.

### Construction of scaffolds

After the repeats identification and the graph simplification steps, most of the contigs were connected in linear stretches on the assembly graph. There were, however, some complex regions that required addition manipulation. We referred to a contig as a divergent node if it linked more than two nodes in the graph (Figure 3). We searched for unique nodes at the end of this complex region and got through this region if there were any long reads that joined two unique nodes. Otherwise, we stopped travelling the graph in the forward direction and switched to the reverse direction. Similarly, the search along the reverse direction of the graph stopped at the end of a linear stretch or at a divergent node. The process was then repeated using an unvisited node as the starting node. The procedure ended after traversing all the unvisited and unique nodes in the graph and outputted all linear paths. Finally, the gap-size between contigs was calculated. If the gap-size value was negative, the contigs were merged into a combined contig, and if the value was positive, a gap was inserted between the contigs (a gap was represented by one or more undefined ‘N’ nucleotides, depending on gap-size).

**Figure 3.**
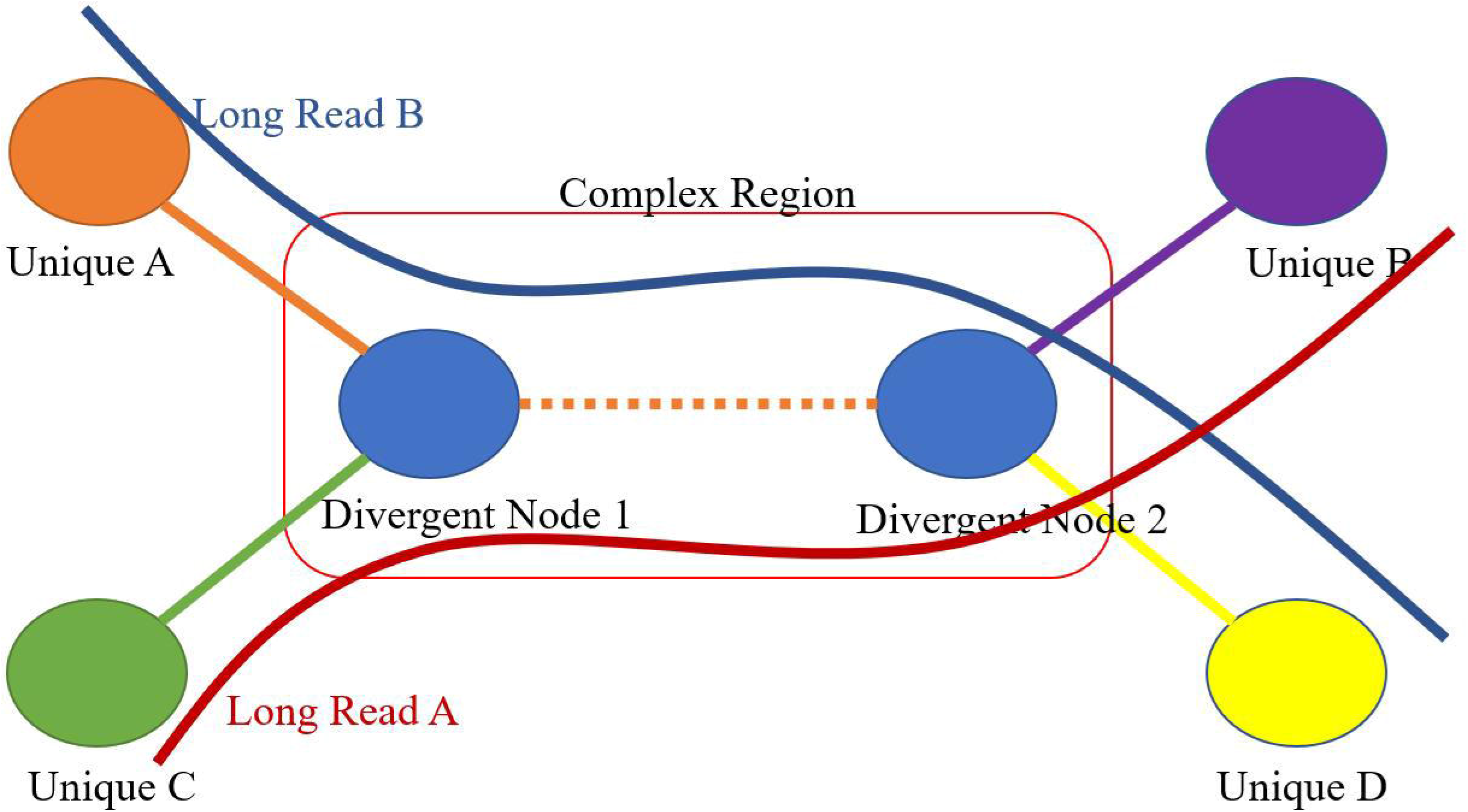
The schematic illustration for travelling complex region.

### Datasets

All tested data were downloaded from published and released datasets (See Table 1). The NGS data of *E. coli* (EAR000206) and *S. cerevisiae* (SRR527545 and SRR527546) were downloaded from EBI and NCBI, respectively, where the NGS data of *D. melanogaster* and *O. sativa* were simulated from their latest reference genome using pIRS (version 1.11) with parameters -x 50 and -c 0, respectively. The SMRT long reads datasets for 5 organisms were published by PacBio^®^: SRX669475 and SRX533603 for *E. coli*, SRX533604 for *S. cerevisiae*, SRX499318 for *D. melanogaster*, SRR3743363 for *O. sativa*, and SAMN02744161 for *H. sapiens* (CHM1). We selected the first 20-fold coverage of each SMRT dataset for comprehensively assessing all scaffolders and we chose 3 different coverages, *i.e*. 1, 5 and 10 -fold, for 2 small genomes (*E. coli* and *S. cerevisiae*) to test all scaffolders performances on lower depths. For the long reads of the ONT dataset, datasets were referred to LINKS and *H. sapiens* (NA12878) with ONT-FULL (ERX708228) for *E. coli*, ONT-ALL (ERX708228) for *E. coli*, ONT-RAW (ERX708228) for *E. coli*, ONT-NANOCORR (SRP055987) for *S. cerevisiae*, ONT-RAW (SRP055987) for *S. cerevisiae* and PRJEB23027 for *H. sapiens*, respectively.

### Draft assembly procedure

The draft genomes for *E. coli*, *S. cerevisiae*, *D. melanogaster* and *O. sativa* were constructed using SOAPdenovo2 taking genome size and contiguity into account. We use two subroutines for *E. coli:* 1) pregraph with −k 51 and −R parameters and 2) contig with −R parameter. We used two similar subroutines for *S. cerevisiae:* 1) pregraph with −k 29 and −R parameters and 2) contig with −R parameter. The draft assemblies for *D. melanogaster* was also constructed using two subroutines: 1) pregraph with −k 61 and −R parameters and 2) contig with −R parameter. For *O. sativa*, we used the subroutine all with −K 63 −p 24 −d 1 −R −F. The two small genomes (*E. coli* and *S. cerevisiae*) were also assembled by SPAdes with the “careful” parameter. To assess the performances between LINKS and the other scaffolders on the ONT long read, the draft assemblies for *E. coli* and *S. cerevisiae* were referred to LINKS. The *H. sapiens* CHM1 and NA12878 draft assemblies were from Steinberg *et al*. [31] and Weisenfeld *et al*. [32]. Table 2 lists the statistics for all of the draft assemblies.

### System

All analysis was performed on a 1 Tb memory Linux machine with 48 CPUs incorporating Hyper-threading technology.

### Source code

LRScaf is written in Java™ and is capable of running on all platforms including Linux, Windows, and Mac if Java Running Environment (JRE) was installed. The source code is available on GitHub (https://github.com/shingocat/lrscaf). We provide a packaged jar file which could be used straight out of the box and the compilation steps for advanced users.

## Additional file

Additional file 1: Long Reads (< 30 Kb) Distribution of Pacific Biosciences^®^ SMRT, and Additional file 2: Long Reads (<30 Kb) Distribution of Oxford Nanopore Technologies^®^ nanopore.

## List of abbreviations

BLASR: Basic Local Alignment with Successive Refinement;
NGS: Next Generation Sequencing;
TGS: Third Generation Sequencing;
SMRT: Single Molecule Real Time;
ONT: Oxford Nanopore Technologies; LRScaf: Long Reads Scaffolder

## Declarations

### Ethics approval and consent to participate

Not applicable.

### Consent for publication

Not applicable.

### Availability of data and materials

The datasets generated and/or analyzed in our study are available in the NCBI repository with accession number listed in Table 1. The datasets synthesized using pIRS are available from the corresponding author on request.

### Competing interests

The authors declare that they have no competing interests.

### Funding

This research was funded by the Dapeng New District Special Fund for Industrial Development [KY20160204, KY20150113]; and the National Key Research and Development Program of China [2016YFC1200600]; and the National Natural Science Foundation of China [31571353]; and the Fundamental Research Funds for Central Non-profit Scientific Institution [Y2016PT54]; and the Fund of Key Laboratory of Shenzhen [ZDSYS20141118170111640]; and the Agricultural Science and Technology Innovation Program; and the Shenzhen Science and Technology Research Funding [JSGG20160429104101251]; and the Key Forestry Public Welfare Project [201504105]; and the Agricultural Science and Technology Innovation Program Cooperation and Innovation Mission [CAAS-XTCX2016].

### Authors’ Contributions

MQ conceived and implemented the method. MQ, ALL, FLZ and HF analyze SMRT dataset characters. MQ, SGW, and LLD analyze ONT dataset characters. MQ wrote the article. YXC and JR supervised the study. All authors read and approved the final manuscript.

## Acknowledgments

We would like to thank anonymous reviewers for their comments in revising the manuscript.

## Author details

Genome Analysis Laboratory of the Ministry of Agriculture, Agricultural Genomics Institute at Shenzhen, Chinese Academy of Agricultural Sciences, No. 7, Pengfei Road, Dapeng District, Shenzhen, China.

---

Supplementary Table 1. The performances for *E. coli* and *S. cerevisiae* based on draft assemblies generated by SOAPdenovo2 and SPAdes using 1, 5, and 10 -fold coverages of PacBio long reads.

Supplementary Table 2. The performances for *E. coli* and *S. cerevisiae* based on draft assemblies generated by SOAPdenovo2 using 20-fold coverage of PacBio long reads.

Additional file 1: Pacific Biosciences SMRT long reads (< 30 Kb) distribution.

Additional file 2: Oxford Nanopore Technologies long reads (< 30 Kb) distribution.

